# Chromatin structure from high resolution microscopy: scaling laws and microphase separation

**DOI:** 10.1101/2023.08.01.551471

**Authors:** Loucif Remini, Midas Segers, John Palmeri, Jean-Charles Walter, Andrea Parmeggiani, Enrico Carlon

## Abstract

Recent advances in experimental fluorescence microscopy allow high accuracy determination (resolution of 50 *nm*) of the 3D physical location of multiple (up to ∼ 10^2^) tagged regions of the chromosome. We investigate publicly available microscopy data for two loci of the human Chr.21 obtained from multiplexed FISH methods for different cell lines and treatments. Inspired by polymer physics models, our analysis centers around distance distributions between different tags, aiming to unravel the chromatin conformational arrangements. We show that for any specific genomic site, there are (at least) two different conformational arrangements of chromatin, implying coexisting distinct topologies which we refer to as phase *α* and phase *β*. These two phases show different scaling behaviors: the former is consistent with a crumpled globule while the latter indicates a confined, but more extended conformation, as a looped domain. The identification of these distinct phases sheds light on the coexistence of multiple chromatin topologies and provides insights into the effects of cellular context and/or treatments on chromatin structure.

## I. INTRODUCTION

Understanding the organization of eukaryotic chromosomes is an issue of broad interest and intensively studied both experimentally [1–8] and computationally [9– 17]. This interest stems from the fact that the 3D chromatin structure has a strong influence on several genomic processes such as transcription or replication [18, 19]. Long DNA molecules go through high compaction when chromosomes condense during mitosis. During the interphase, genes are actively transcribed and chromosomes are unpacked and distributed throughout the cell nucleus but still possess a remarkable level of spatial organization. This organization is hierarchical, involving different structures at different genomic length scales. Starting from whole chromosome lengths (∼100 Mb) down to ∼ 100 Kb the DNA organization can be summarized as follows: 1) During the interphase, different chromosomes do not mix but occupy well-defined territories inside the cell nucleus. 2) Each chromosome is split into several stretches of gene-rich domains (euchromatin) alternating with gene-poor domains (heterochromatin). Euchromatin tends to be more open than heterochromatin. Moreover stretches of the same chromatin species aggregate forming so-called A and B compartments. 3) At scales of ∼1 Mb and below the chromatin is assembled in Topologically Associating Domains (TADs), which are region where chromatin interacts more frequently.

The biological function of TADs is still debated as well as the mechanisms leading to their origin. A prevalent line of thoughts suggests that TADs originate from a loop extrusion mechanism [20]. Other models of genome organization have been proposed in the literature such as the strings and binders model [11], the block-copolymer model [21] or the diffusing transcription factor model [12]. It is very likely that different mechanisms operate simultaneously to contribute to the complex hierarchical organization of eukaryotic chromatin.

Advanced techniques such as Chromosome Conformational Capture (3C) [1], particularly its high-throughput sequencing (Hi-C) version [2], have played a central role in unveiling the chromatin organization. Hi-C provides genome-wide data for contact probabilities between pairs of genomic sites and was instrumental to understand the folding of chromosomes at different length scales. Although it is by no doubt a very powerful technique, Hi-C has also some limitations. For instance, it provides indirect information on physical distances (via contact probabilities), it typically requires the averaging over a large number of chromosomes (∼10^6^) and may contain some systematic biases [22], therefore raw data require normalization and other pre-processing steps. Fluorescence in situ hybridization (FISH, Fig. 1) has also been used to investigate chromosomal structure [23]. FISH uses fluorescent probes that are complementary to desired chromosomal regions. After hybridization, the 3D location of the probes can be determined from fluorescence microscopy. For a long time FISH remained a very low throughput technique, the main issue being the limited fluorescent probes with different emission spectra available so that only a few sites can be visualized. However, quite recently, a multiplexed FISH (m-FISH) method was developed [4]. Using unique readout sequences and sequential hybridization cycles, the method allows to determine the positions of *N ∼* 50 *−* 100 chromosomal sites with high resolution (*<* 50 nm), Fig. 1. Measurements are done on samples containing up to *M ∼* 10^4^ distinct chromosomes. Although, unlike Hi-C, it is not a genome-wide technique, m-FISH data reproduce well the Hi-C contact probabilities for the same genomic regions [4]. In addition, m-FISH reveals cooperative (i.e. many-body) chromatin interactions which cannot be inferred from Hi-C pairwise contact probabilities.

**FIG. 1.**
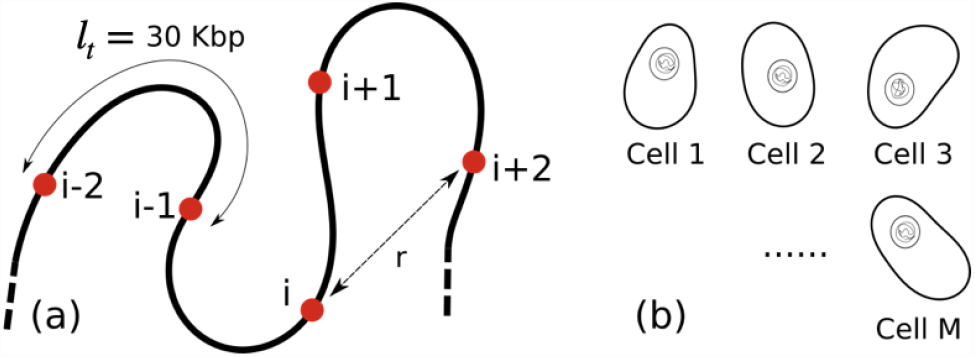
Setup of multiplexd FISH as developed in Ref. [4]. (a) 1 ≤ *i* ≤*N* genomic sites equally spaced are targeted by FISH probes during sequential hybridization cycles. In the setup of [4] the genomic distance between consecutive tags was *l*_*t*_ = 30 Kbp. The 3D spatial coordinates of the tagged sites are then read by high-resolution fluorescence microscopy. (b) The experiment is repeated over *M* distinct copies of chromosomes. The values of *N* and *M* for the experimental data used in this paper are given in Table I. We analyzed here the distance distributions for any two given sites *i* and *j*.

The aim of this paper is to analyze a set of m-FISH data focusing on distance distributions of any two pairs of labels *i* and *j*. In particular, we analyze histograms *P*_*ij*_(*r*) of distances between the tags *i* and *j*. The large number of distinct samples (*M* ∼ 10^4^) implies good statistics, therefore these histograms can be accurately determined. Our analysis reveals that there are (at least) two different modes of chromatin organization at any given genomic site. We report scaling laws characterizing the growth of the chromatin domains when increasing the genomic distance. Overall, our work presents a polymer physics-inspired analysis to characterize chromatin structure. Importantly, this method is straightforward to be implemented and applied to “raw” experimental data.

The paper is organized as follows. Section II presents the method we propose based on the analysis of the experimental distance distribution histograms. Section III focuses on the analysis of the extracted scaling behaviour of typical configuration radii from different cell lines. Section IV discusses, using polymer physics models, the emerging regularities from typical chromatin conformations from the experimental data. Appendices provide technical details on the polymer models considered and a full analysis of the whole dataset histograms.

## II. DISTANCE DISTRIBUTION HISTOGRAMS

Table I summarizes the main features of the four data sets from [4] analyzed in this paper. Each dataset corresponds to a cell line whose properties will be detailed later in Section III. These experiments target two different loci of human chr. 21: a 2 Mbregion (Chr21:28Mb-30Mb) labeled by *N* = 65 tags and a 2.5 Mbregion (Chr21:34.6Mb-37.1Mb) labeled by *N* = 83 tags. These regions contain several TADs across multiple cell lines [4]. More data are available in [4], but these 4 sets were selected because they contain a large number of independent measurements, ranging from *M* = 4871 to *M* = 13997, (Table I) from which accurate histograms are obtained. The histograms from the *M* experimental samples are normalized as follows

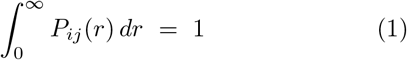

For an ideal polymer, the distance distribution is a gaussian in the vector distance 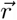, while for its length 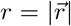 takes the form

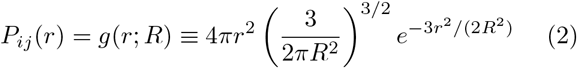

where the term 4*πr*^2^ comes from the jacobian of the distribution. The above distribution is normalized as (1) and contains the mean-squared radius 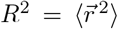 as a single parameter. Despite its simplicity the gaussian chain model is a reasonable approximation to complex phases of polymeric matter, such as polymer melts. A well-known result of polymer physics is that a single test chain immersed in a melt of other chains behaves as an ideal polymer [24]. This is because self-avoidance gets screened by the surrounding polymers. The mean square radius of an ideal polymer scales as

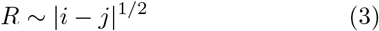

where the exponent 1*/*2 describes the universal scaling behavior of random walks. It has been recently shown [25] that the introduction of suitable monomer-monomer harmonic pairwise interactions in a bead and spring polymer model leads to an equilibrium distance distribution as (2), but with a scaling described by

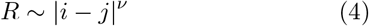

By tuning the monomer-monomer interactions one can generate any values of the exponent in the interval 1*/*3 ≤ *ν* ≤ 1*/*2 [25]. The statistical properties of the model are those of a fractional Brownian motion (fBm), which was invoked as a model for chromosome arrangement in cells [25], as well as in other problems of polymer dynamics [26, 27]. We note that the case *ν* = 1*/*3 is the exponent of a crumpled globule [28], a phase discussed in some more detail in Sec. IV. Therefore, the gaussian distribution (2) encompasses a broad range of polymer models.

**TABLE I.** Summary of experimental data analyzed from [4]. These target two different regions of chr.21: a 1.2 Mb region in the 28 − 30 Mb genomic position and another one 2.5 Mb in the 34 − 37 Mb position. Consecutive tags are separated by a genomic distance of 30 kbp. HCT116 is a colon cancer cell line, IMR90 a lung fibroblast cell line and K562 an erythroleukemia cell line.

| n. | Experiment/Cell line | $N$ | $M$ |
| --- | --- | --- | --- |
| 1 | HCT116_chr21-34-37Mb_untreated | 83 | 11631 |
| 2 | HCT116_chr21-34-37Mb_6h-auxin | 83 | 9526 |
| 3 | IMR90_chr21-28-30Mb | 65 | 4871 |
| 4 | K562_chr21-28-30Mb | 65 | 13997 |

Figures 2 (HCT116 and auxin-treated HCT116 cell lines) and 3 (IMR90 and K562 cell lines) show several plots of experimentally measured histograms *P*_*ij*_(*r*) (circles), where the tags *i* and *j* considered are reported in each graph and the horizontal scale is the spatial distance in micrometers (*μm*). The green solid lines are a one parameter fit to Eq. (2), where the mean-squared radius *R* is the only fitting parameter. We note that the model does not fit well the experimental data and the deviations are particularly strong for close tags, e.g. small |*i* − *j* | (say within range of |*i* − *j* | ≈ 4 corresponding to a genomic distance of 120 Kbp). As the genomic distance increases, i.e. larger |*I* − *j*|, the experimental distributions gets closer to (2), seen by the trend to merging of the green line (gaussian fit) to the black circles (experiments) in Figs. 2 and 3. This merging trend is however stronger in the untreated HCT116 and the IMR90 lines, while there remains a substantial gap between the single and double gaussian fits for large genomic distances | *i* − *j* | = 25 for the auxin-treated HCT116 and the K562 lines.

**FIG. 2.**
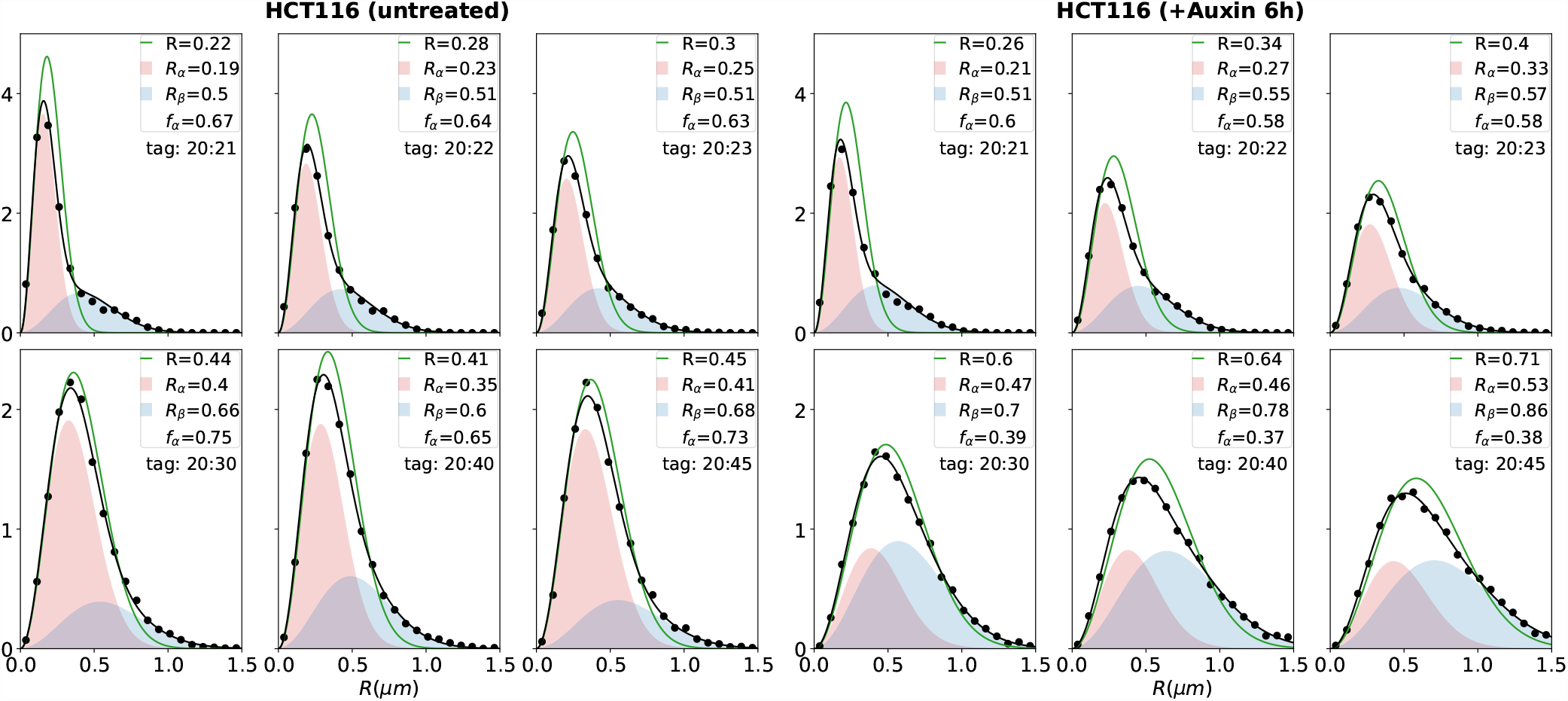
Histogram of distance distributions *P*_*ij*_ (*r*) from experimental data (black circles) for different tag locations *i, j* for experiments of the HCT116 cell line (sets n.1 and n.2 of Table I). The values for *i* and *j* are indicated in each graph. In the two sets, we fix *i* = 20. The tag *j* is increased from *j* = 21 (top left) to *j* = 45 (bottom right). Solid lines are fit to a single gaussian model (Eq. (2), green line) and double gaussian model (Eq. (5), black line). The legends in each plot show the value of *R* for a single gaussian model (Eq. (2), green) and of *R*_*α*_, *R*_*β*_ and *f*_*α*_ for the double gaussian model (Eq. (5)) The red and blue filled areas denote the contribution of the two components of the fit. The distance in the horizontal axis is in *μ*m.

**FIG. 3.**
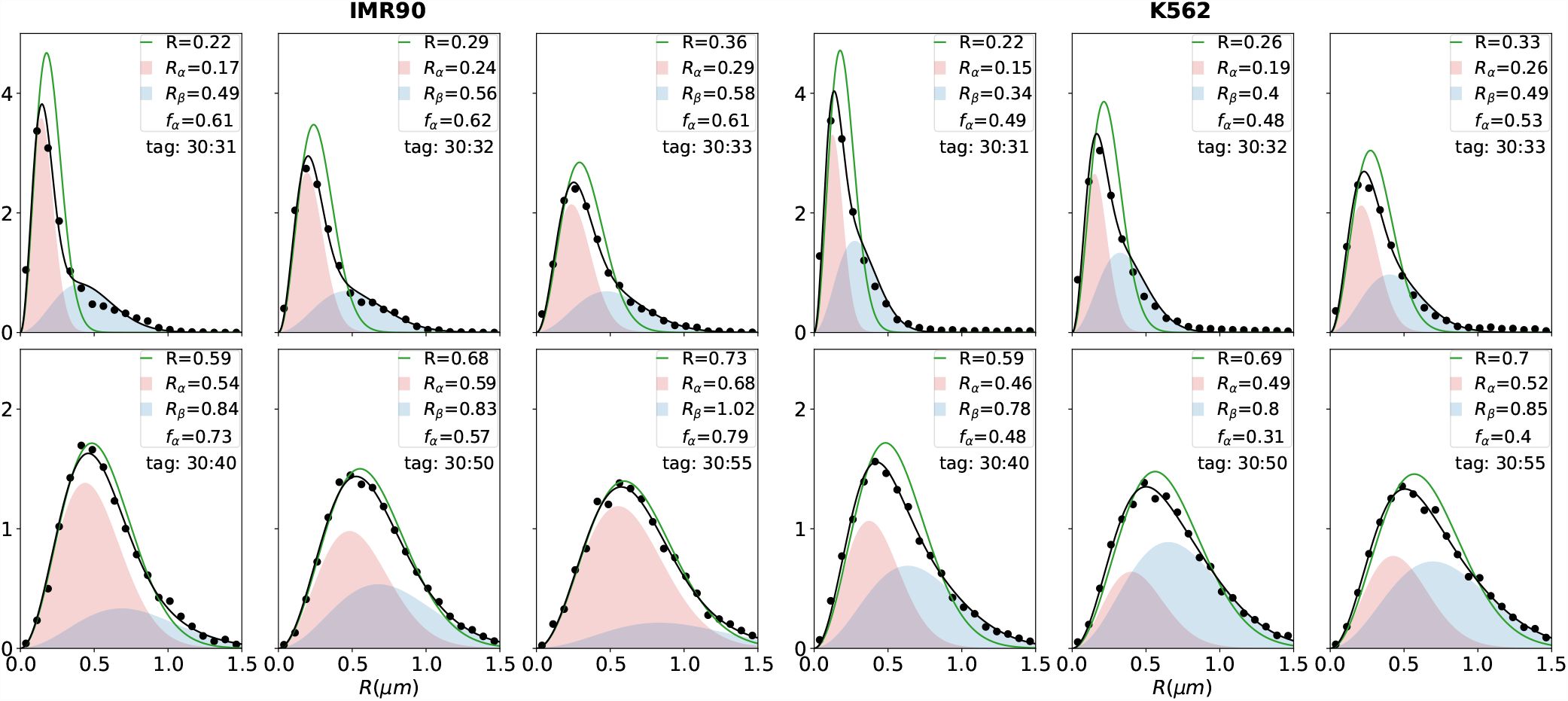
Same analysis as in Fig. 2 for experiments n.3 (IMR90 cell line) and n.4 (K562 cell line) of Table I.

The histograms for small |*i* − *j*| suggest that experiments describe two coexisting populations (phases), therefore we fitted the data using:

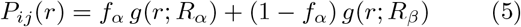

where the parameter 0 ≤*f*_*α*_ ≤1 (respectively *f*_*β*_ = 1 − *f*_*α*_) is the fraction of chromosomes with mean radius *R*_*α*_ (respectively *R*_*β*_) around the tags *i* and *j*. We will refer to the two phases as *α* and *β*. The two-phase model (5) provides an excellent fit for the experimental data. These fits are shown as solid black lines in Figs. 2 and 3. The colored areas are the contributions of the two gaussian components to the final fit. The two-phase model is fitted to the data using *R*_*α*_, *R*_*β*_ and *f*_*α*_ as adjustable parameters. These three values are reported in the legends of Figs. 2 and 3. The radii change with changing *i* and *j* and, as expected, increase for increasing |*i* −*j*|. The growth rate is however different for *R*_*α*_ and *R*_*β*_, as discussed in the next Section.

## III. SCALING BEHAVIOR OF *R*_*α*_ AND *R*_*β*_

Figures 4 and 5 show the extracted scaling behavior for the four different cell lines. These are log-log plots of the mean radii *R*_*α*_ and *R*_*β*_ vs the genomic distance *l*_*t*_ |*i−j*| for fixed *i* and varying *j*, where *l*_*t*_ = 30 Kb is the spacing in base pairs between two consecutive tags of the experiments (see Fig. 1). The horizontal axis is thus the genomic distance measured in bp, which extends up to about 2.5 Mbp. We characterized the scaling behavior of the radii fitting *R*_*α*_ and *R*_*β*_ to (4) for close tags, i.e. |*i* −*j*| ≤6 corresponding to a maximal genomic distance of 180 Kb. This choice is dictated by two factors: (*i*) we observe that the scaling of *R*_*α*_ and *R*_*β*_ follows an approximate power-law behavior for a limited distance between tags and (*ii*) the values of *R*_*α*_ and *R*_*β*_ are more accurately determined from fitting histograms for small |*i* −*j*| as well. This is because there is typically a bigger gap between *R*_*α*_ and *R*_*β*_ for small |*i* −*j*|, therefore the two radii can be more reliably extracted from the data analysis. Table II gives a summary of the average exponents for the four different cell lines averaged over all the tags. The exponents give a measure of the average geometrical properties of the two phases. With the exception of the K562 cell line, we find consistently *ν*_*α*_ *> ν*_*β*_. The properties of the different cell lines will be discussed in the following.

**TABLE II.** Mean values of the geometric exponents *ν*_*α*_ and *ν*_*β*_ for the different data sets. The last column gives the value of *f*_*α*_ averaged on all data in the range |*i − j*| *≤* 5.

| Data set | $\nu_\alpha$ | $\nu_\beta$ | $f_\alpha$ |
| --- | --- | --- | --- |
| HCT116 (untreated) | 0.35(10) | 0.11(7) | 0.64(5) |
| HCT116 (+Auxin 6h) | 0.40(6) | 0.10(3) | 0.60(5) |
| IMR90 | 0.31(11) | 0.11(6) | 0.61(8) |
| K562 | 0.38(8) | 0.35(7) | 0.53(8) |

**FIG. 4.**
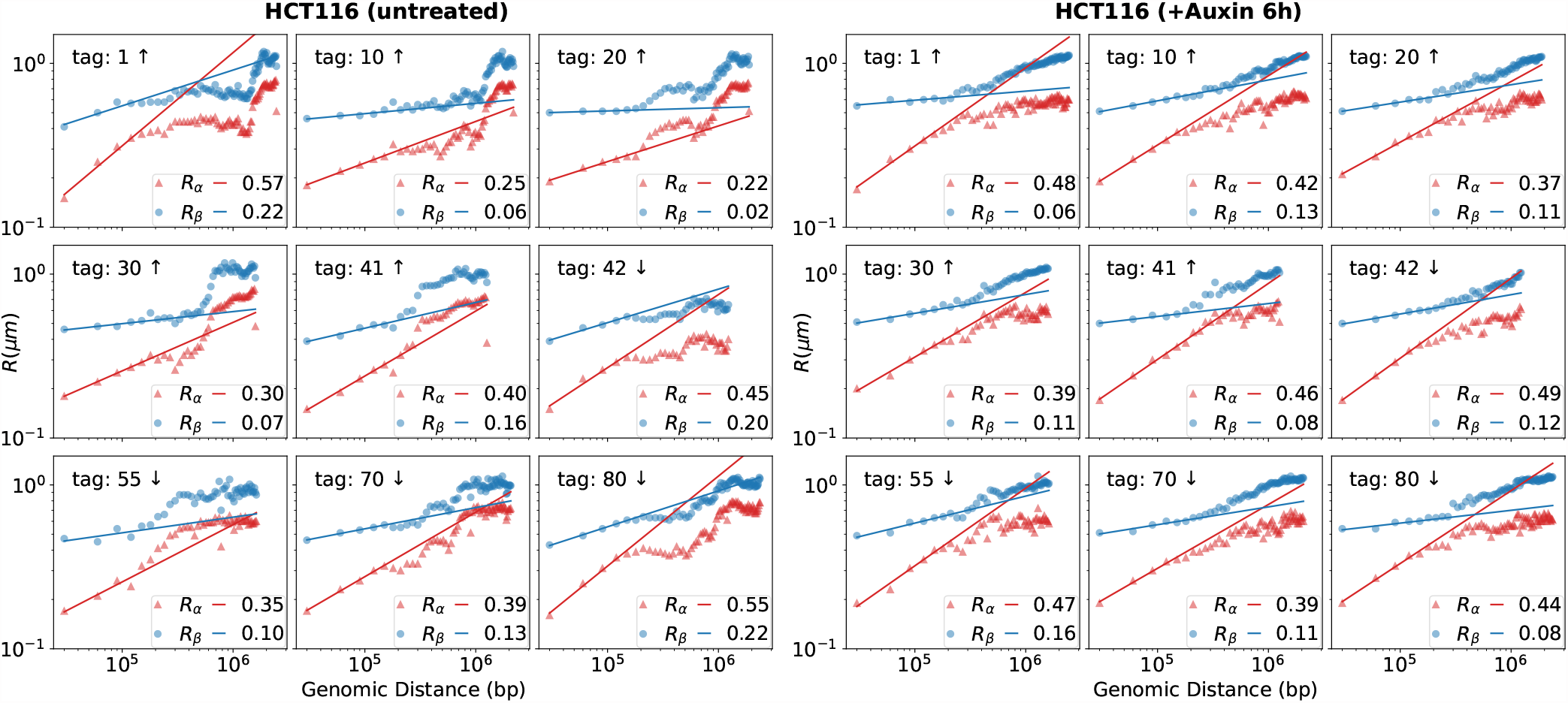
Log-log plots of scaling behavior *R*_*α*_ (red triangles) and *R*_*β*_ (blue circles) vs. genomic distance for some selected tags of the HCT116 cell. Left: untreated and Right: 6h-auxin treated cells. The reference tag used is given in each plot. The notation “tag: 1 *↑*“means that *R*_*α*_ and *R*_*β*_ are calculated for *i* = 1 and *j* = 2, 3 … *N* and “tag: 55 *↓*“plots the data for *i* = 55 and *j* = 54, 53, … 1. The solid lines are power-law fits over the first 5 point of each set.

**FIG. 5.**
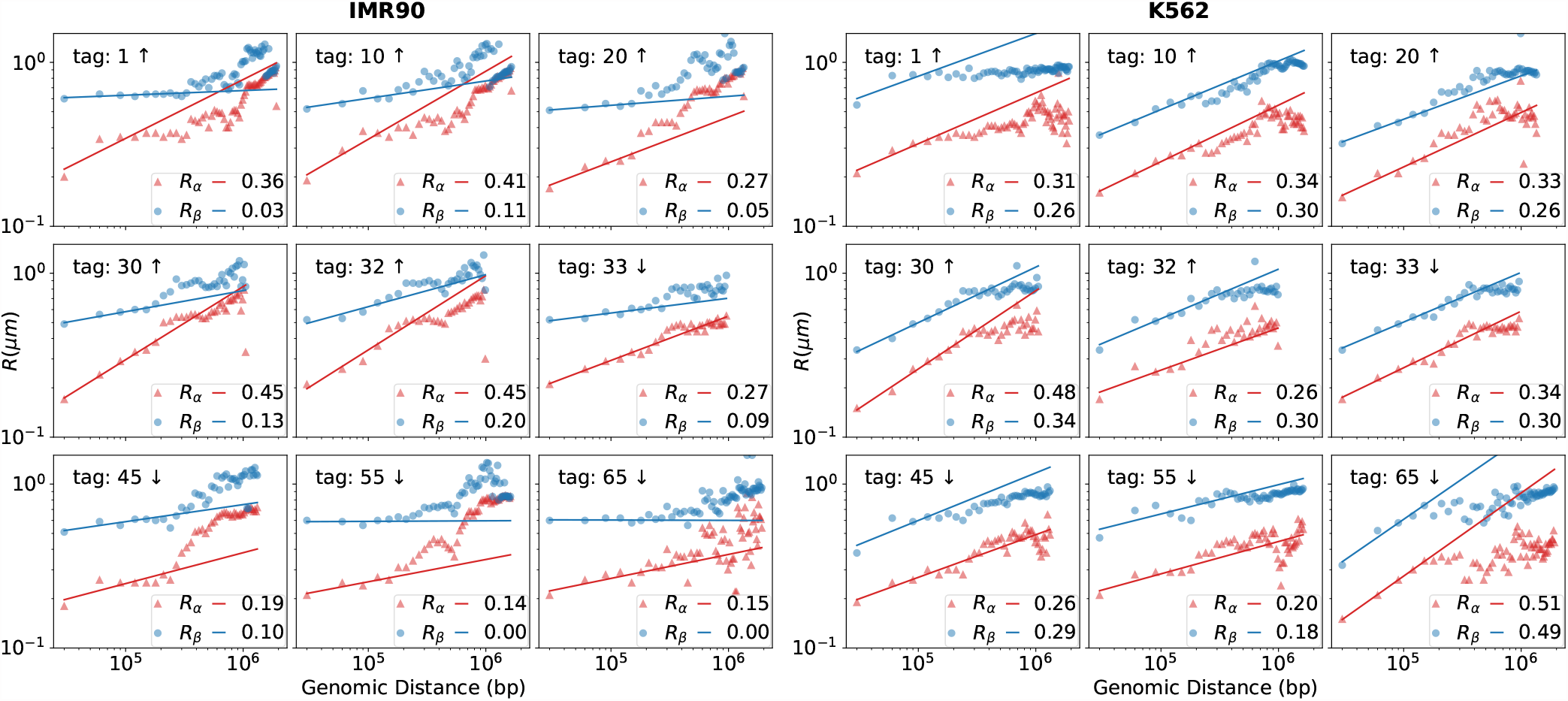
Same analysis as in Fig. 4 for the cell lines IMR90 (left) and K562 (right).

### A. HCT116 cell line

HCT116 are human colon carcinoma cells. Figure 4 shows the scaling behavior of *R*_*α*_ and *R*_*β*_ for a few selected tags of the untreated (left) and the auxin-treated (right) cases for HCT116. Auxin degrades cohesin and it is therefore expected to affect the TAD structure. We find, however, very little difference between the exponents in the two cases, apart from a possibly slightly higher *ν*_*α*_ in the auxin-treated case, see Table II. This small increase is a signature that the *α*-phase is slightly less compact in the auxin-treated case. This is consistent with the fact that by degrading cohesin, auxin releases some constraints. The overall little difference in exponents in the two cases agrees with the conclusions of Bintu et al. [4]. The authors found that the *ensemble averaged*, contact matrices from mFISH show no sign of TADs in the auxin-treated cells. However, TADs were still visible at the *single cell* level. As a matter of fact, as explained in [4], while TAD boundaries in the untreated HCT116 cells are pinned at specific genomic sites, in the auxin-treated line, they are found in different genomic positions for each cell, therefore washing out any TAD signature in the ensemble average as done in Hi-C data analysis. The effect of pinned vs. variable boundaries is also visible in Fig. 4: the scaling of *R*_*α*_ and *R*_*β*_ is smoother and power-laws extend to a broader interval in the auxin-treated cells as compared to the untreated ones. In the latter, several “bumps”, which are likely the effect of strongly pinned domains, are visible.

### B. IMR90 cell line

Figure 5 (left) shows the scaling properties of the IMR90 lung fibroblast cell line. The exponents *ν*_*α*_ and *ν*_*β*_ are consistent with those observed in the untreated HCT116 cell lines, see Table II. As in the untreated HCT116 case, one observes from the irregular shape the effects of TADs pinning at specific sites. This is consistent with ensemble-averaged contact map results [4].

### C. K562 cell line

The last case analyzed was the K562 erythroleukemia cell line, shown in Fig. 5 (right). Unlike the previous cases, we observe for K562 quite close exponents for the two phases *ν*_*α*_ ≈ *ν*_*β*_, thus *ν*_*β*_ is substantially larger than in the other cell lines, see Table II. This suggests a different type of chromatin organization for K562 as compared to HCT116 and IMR90. The origin of the peculiar behavior is presently not understood. On view of the numerical value of *ν*_*β*_ it is more logical to characterize this cell line as having two phases of type *α* but with two distinct values of *R*_*α*_.

## IV. DISCUSSION

In this paper, we have analyzed mFISH data from the experiments reported in Ref. [4] for human chr.21 for different cell lines. Our main result is the evidence of two different coexisting phases of chromatin, which we referred to the *α* and *β* phases. The range of genomic length explored by these experiments goes from 30 Kbp to 2.5 Mbp, but the characteristic radii *R*_*α*_ and *R*_*β*_ Can be confidently obtained from data for distances up to approximately 0.5 Mbp. Therefore our claims are limited to this range of lengths, which is roughly the genomic distance at which TADs are observed. We characterized the two phases using scaling exponents *ν*_*α*_ and *ν*_*β*_. Interestingly, these exponents are different in the two phases (except for the K562 cell line) indicating substantially different spatial organizations, which we discuss in this Section.

### A. Phase *α*

The exponent we extracted for the phase *α* is close to that expected for a crumpled globule, which is *ν* = 1*/*3 [28–30]. The crumpled globule is a metastable phase arising from the rapid condensation of a self-attracting polymer following a temperature quench. The polymer condenses, but it has no sufficient time to relax to a true equilibrium conformation. The rapid condensation prevents the end points of the long polymer to retract and to form knots, so the crumpled globule remains unentangled [30]. This is different from an equilibrated compact phase, which is strongly entangled and forms knots. The absence of knots makes the unfolding of crumpled globules much more rapid than that of equilibrated conformations [31]. Crumpled globule polymeric phases have been discussed in melts of polymer rings [9] and are believed to be relevant for genome folding structures [31, 32]. Figure 6(a) illustrates the typical conformation of a crumped globule. Nearby chromatin segments (indicated with different colors in Fig. 6(a)) tend to condense and the process continues hierarchically involving longer and longer length scales, with the end-points playing no specific role in the process.

**FIG. 6.**
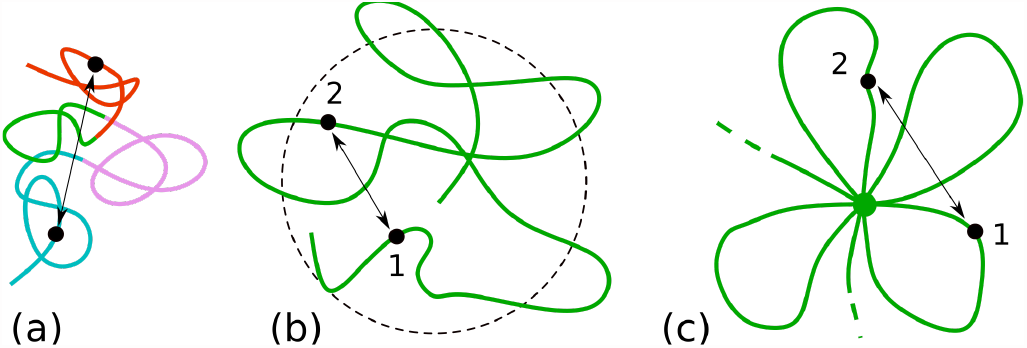
Possible phases of chromatin: (a) crumpled globule, (b) confining due to external potential, and (c) flower. (a) is a candidate for the phase *α* and (b,c) for the phase *β*. In (a) different segments of the chromatin fiber are colored differently to illustrate the geometry of the globule. The rapid condensation leads to collapses at the local scale involving vicinal regions. In (b) and (c) strong confinement implies *R*_*β*_ *∼ s*^*0*^ for *s* sufficiently long.

### B. Phase *β*

The phase *β* is characterized by a very small exponent, which is an indication of strong confinement. Figure 6(b) and (c) illustrate two possible mechanisms of polymer confinement. In (b) the confinement is due to an external potential forcing the monomers to remain within a certain range from a fixed origin. This could be due to the formation of a droplet by, for instance, liquid-liquid phase separation, with chromatin preferentially absorbed by the droplet. An alternative interpretation is a flowerlike conformation (c) where loops, possibly forming via an extrusion mechanism, are bound to a central hub. For both types of conformations (b) and (c) one has *R*_*β*_ ∼ *s*^0^, i.e. the characteristic radius becomes independent on the typical genomic length *s* beyond some threshold value *s > s*_0_. Here *s*_0_ is the typical genomic length of the polymer to reach the droplet surface and back to the origin in (b) or the typical loop genomic size in (c). A mathematical model for the case (b) is discussed in Appendix A, but the *s*-independence can also be explained intuitively. Let us consider the point “1” and “2” in Fig. 6(b), which we assume are separated by a genomic distance *s > s*_0_. To reach point “2”, the polymer chain bounces a few times with the boundaries of the confined region. In doing that any information about the length *s* is lost and the probability distribution that the two points have a distance 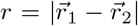 does not depend on *s*. For confinement of type of Fig. 6(c), if the points “1” and “2” are on different petals of the flower-like configuration then again the distance distribution of 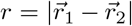 will not depend on the genomic distance *s*. Experimental data show no “ideal” confinement (*ν* = 0) as the average exponent is non-vanishing, although very small *ν*_*β*_ ≈ 0.1. We note that, although confined, the phase *β* is characterized by a radius *R*_*β*_ *> R*_*α*_ in the range of genomic distances up to ≈ 2 *·* 10^5^ bp.

## C. Coexisting topologies

The picture emerging from this analysis reveals different coexisting conformations in which in which the chromosome of each cell alternates between micro-domains of phases *α* and *β*. To illustrate this behavior we plot in Fig. 7 (solid black lines) the local radius of gyration *R*_*g*_(*i*) calculated over five subsequent tags centered around the tag *i*. The plots are obtained from the data of six randomly selected cells for the cell lines 1, 2 and 3 of Table I. While so far we have analyzed histograms over the whole set of *M* distinct cells, here we perform a singlecell analysis where Fig. 7 illustrates a marked cell-to-cell variability. The radius of gyration *R*_*g*_(*i*) defined above describes the local structure of the chromatin in a range of 150 Kbp.

**FIG. 7.**
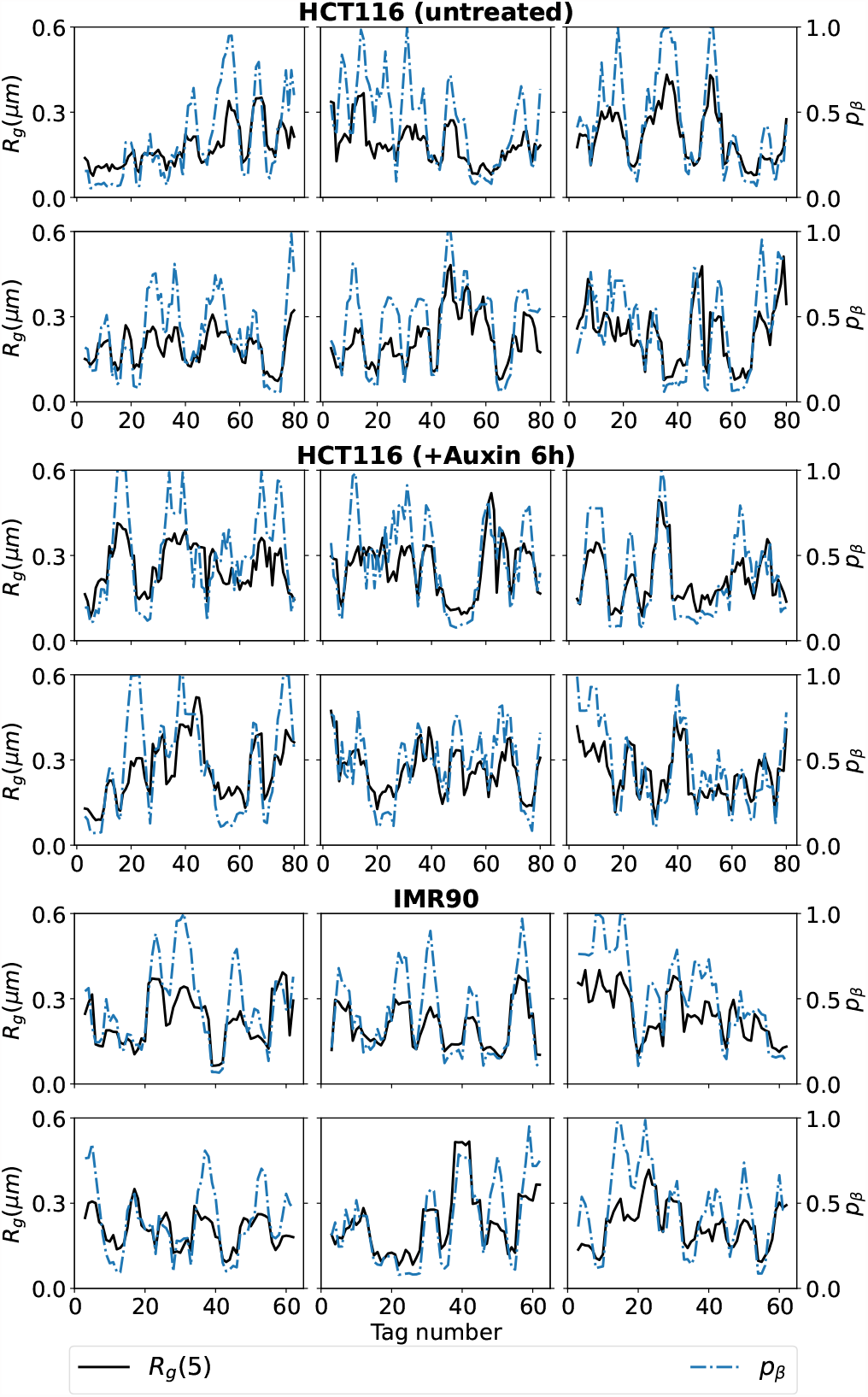
Radius of gyration *R*_*g*_ (solid black) and *β*-phase probability *p*_*β*_ (broken blue) as a function of tag number for different cells and cell lines. *R*_*g*_ is calculated over five subsequent tags. *p*_*β*_ is defined in Eq. (6).

In Fig. 7 one remarks large and small values of *R*_*g*_(*i*) indicative of more open and more compact configurations, as expected for the phases *β* and *α*, respectively.

We also analyzed the *β*-phase probability *p*_*β*_(*i*) as function of tag position *i*, which is defined as follows. We constructed first a global histogram of consecutive tags by averaging *P*_*i,i*+1_(*r*) over all tag positions *i* for all cells within a cell line. We indicate this averaged histogram with ⟨*P*_*i,i*+1_(*r*)⟩_*i*_. From a two gaussian fit (5) of ⟨*P*_*i,i*+1_(*r*)⟩_*i*_ we obtain global values for *f*_*α*_, *R*_*α*_ and *R*_*β*_ over all tags. Given now a measured distance 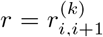 between tags *i* and *i* + 1 for a cell *k* we define the local probability to find the local chromatin conformation in the *β* phase as

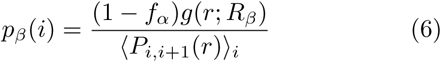

with *g*(*r*; *R*) as in (2). In Fig. 7 we superpose the plots of the local gyration radius *R*_*g*_(*i*) (black solid line) and the local probability *p*_*β*_(*i*) (blue dashed lines) averaged over 5 tags centered around the tag *i*. We thus compare the spatial variability (expressed in tag number) of these physical quantities. Interestingly, the figure shows that domains with a high *β*-phase content co-localize with maxima in the local radius of gyration, whereas clusters with much lower *p*_*β*_ are centered around the lower values of *R*_*g*_ thus with compact regions of the chromatin. The location of these different domains vary strongly from cell to cell, however, the variation of these two quantities is remarkably correlated.

The existence of cooperative interactions, which is an indication of distinct phase behavior, was also pointed out in the single-cell analysis of mFISH data by Bintu et al [4]. In addition, stochastic simulations of a polymer-based chromatin model revealed a heterogeneous folding pattern [33]. This computer model combines several mechanisms of chromosome organization and genomic information. Stochastic simulations from open conformations showed that chromatin forms distinct topologies rather than folding into a single structure [34]. This result is consistent with our findings and the conclusion of this paper. One of the advantages of our analysis is that it captures the heterogeneity of chromatin structures in a simple way from histograms using general concepts of polymer physics such as scaling laws and exponents. Overall, the analysis indicates that the chromatin organization is described by two underlying fractional Brownian motions. In the future, it would be interesting to extend this methodology to more cell lines and also explore chromosomal regions below the 30 Kbp resolution.

## ACKNOWLEDGEMENT

Discussions with M. Barbi, T. Foldes, M. Liefsoens, M. Nollmann are gratefully acknowledged. EC wishes to thank the Laboratoire Charles Coulomb of the University of Montpellier, where part of this work was done, for kind hospitality. This project was supported by the LabEx NUMEV (ANR-10-LABX-0020) within the I-Site MUSE (ANR-16-IDEX-0006). LR acknowledges doctoral thesis support from the LabMUSE EpiGenMed within the I-Site MUSE (ANR-16-IDEX-0006).

## Appendix A Confined Ideal polymer

We discuss here the effect of confinement in the simpler and analytically tractable case of an ideal polymer. We consider a polymer consisting of *N* monomers of average bond length *b* which is subject to an external radial potential *U*(*r*) acting on all its monomers, as depicted in Fig. 6(b). The potential is attractive forcing the polymer to remain confined in the vicinity of the origin. One can consider different forms for *U*(*r*) as infinite well case

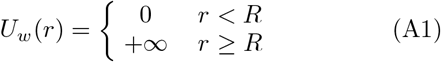

a harmonic potential

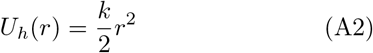

of other types of confinement. The probability distribution for the end point distance 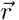 of a segment of length *s* satisfies the differential equation [35]

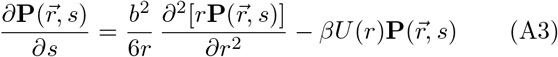

This equation is solved by the separation of variables, e.g. seeking solutions of the type 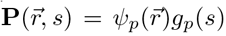, labeled by an index *p*. The general solution is then given by the linear combination

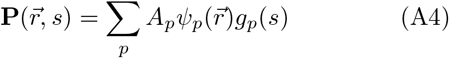

where the coefficients *A*_*p*_ are chosen to satisfy the desired boundary conditions (the full series solution of the infinite-well confinement is given in [36]). Separation of variables leads to a differential equation for *g*_*p*_(*s*) of the type *dg*_*p*_(*s*)*/ds* = −*α*_*p*_*g*_*p*_(*s*) with exponentially decaying solutions 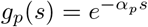. In the limit *s* → ∞ one uses the ground state dominance (GSD) approximation which retains the component with the smallest *α*_*p*_ [37]

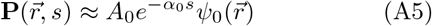

where 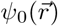 is the ground state solution of the associated Schödinger-like equation

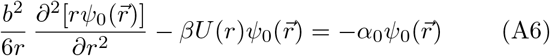

We note that in the GSD approximation the distribution in 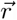 is independent of the polymer length *s*, due to the factorization in (A5). This is different from a free ideal polymer where the probability distribution is a function of the scaled variable 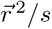.

For a infinite well potential (A1) the solution of (A6) is a spherical Bessel function of order zero

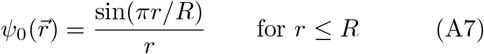

and *ψ*_0_(*r*) = 0 for *r > R*. For a confining harmonic potential (A2) one finds

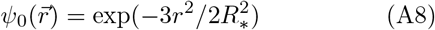

where we have defined 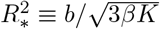. In both cases, we have omitted normalization factors.

We recall that 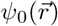, obtained from the ground state dominance (A5), is the probability distribution of the end-point of a very long polymer. The probability that two points along the polymer are separated by a given vector distance 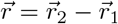 is given by

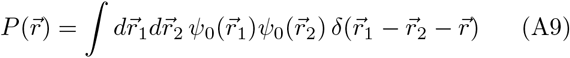

where *δ*() denotes the Dirac delta function. Inserting the following Fourier transform representation of the Dirac delta

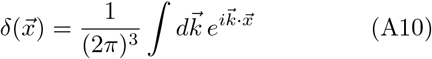

we obtain

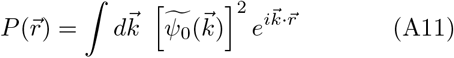

where 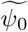 is the Fourier transform of 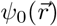. The Fourier transform of a Gaussian function is also a Gaussian, hence the harmonic confinement gives

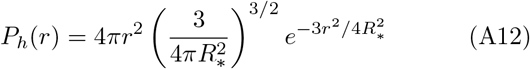

where we have included the jacobian 4*πr*^2^. For the infinite well confinement we solved the problem numerically. We generated independent vector pairs 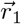 and 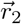 within a sphere of radius *R* distributed according to (A7) and calculated histograms of the distance 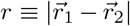, which is shown in Fig. A.1 as solid line. This distribution deviates from a Gaussian one, which is shown as a dashed line. It decays faster than the Gaussian at larger *r* and vanishes for *r >* 2*R* (*r* = 2*R* is the maximal distance between two points within a sphere of radius *R*). For generic confining potential the distribution is not a Gaussian one.

We consider now the case of confinement of an ideal polymer in a flower-like shape conformation as in Fig. 6(c). For simplicity we will assume that each petal has the same length *l* and we number them 1, 2, … *n* according to increasing genomic position. Let us, again for simplicity consider two points in the flower with positions 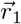,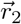 and are located in the middle of petals *i* and *j*. The genomic distance between these points is therefore *s* = *l*| *i* −*j*|. Given 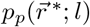 the probability distribution that the mid-point of the petal is at a distance 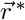 from the center of the flower, the probability distribution that the spatial distance between the two points is equal to 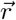 is then given by

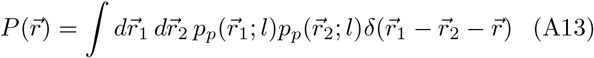

**FIG. A.1.**
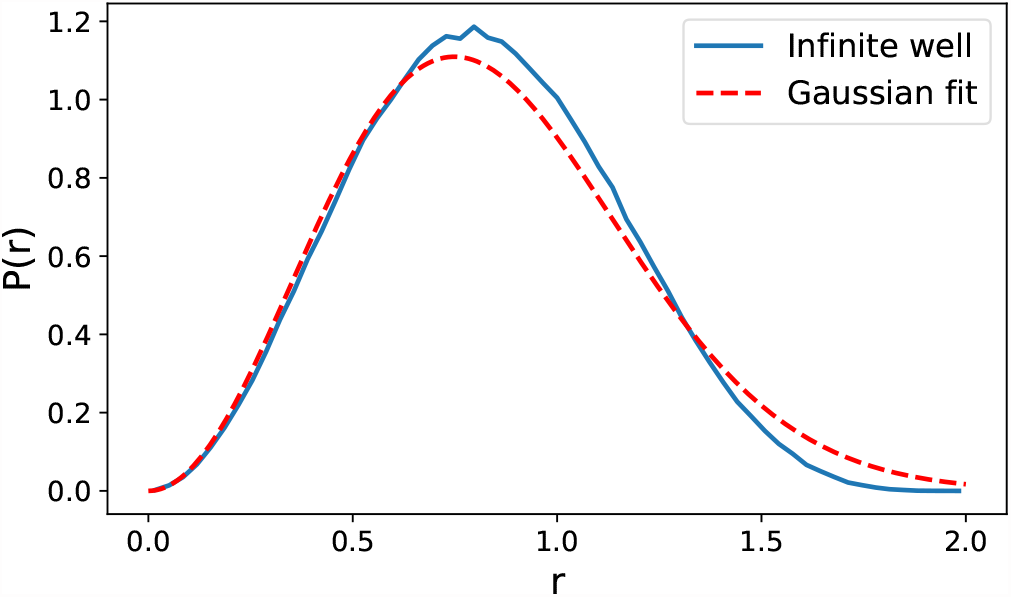
Solid line: Numerical solution of the probability distribution *P* (*r*) of spatial distance *r* between two points separated by a long genomic distance *s* for confinement in the case of infinite well-potential (A1) with *R* = 1. Dashed line: Gaussian fit of *P* (*r*).

**FIG. B.1.**
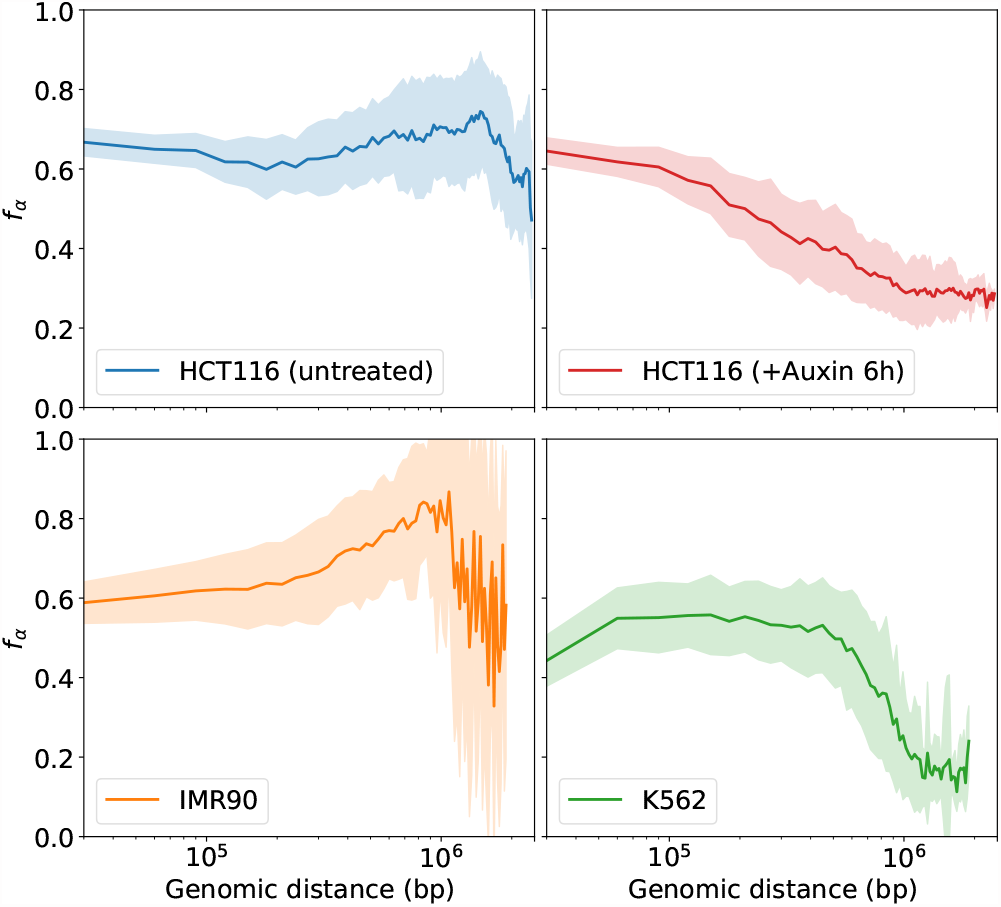
Plot of the average *f*_*α*_, the fraction of the fraction of phase *α*, as a function of the genomic distance (solid colored line). The light colored area indicates the error estimate.

For an ideal polymer 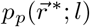 is a Gaussian distribution and so is 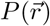 obtained from the previous formula. Most important, this is independent on the genomic distance *s*, hence it predicts a scaling *R* ∼*s*^*ν*^ with *ν* = 0. This is for distances *s > l*, i.e. beyond the length of a petal.

## Appendix B Fractions of *α* and *β* phases

From fits of experimental histograms *P*_*ij*_(*r*) with the two phase model (5) one obtains estimates of *f*_*α*_ and *f*_*β*_ = 1 − *f*_*α*_, the fraction of the two phases for pairs of labels *i* and *j*. Figure B.1 shows the values of *f*_*α*_ (central thick colored lines) averaged over all begin tags *i* as a function of genomic separation *l*_*t*_ |*i* −*j*|, where *l*_*t*_ = 30, 000 is the spacing in base pairs between two consecutive tags of the experiments. The colored areas estimates the variability (standard deviation) over the different data. For a microphase separation we expect a constant value of *f*_*α*_ for a distance range corresponding to the characteristic length of the domains. The data in Fig. B.1 show that *f*_*α*_ ≈ 0.6 for the shortest tags distance |*i* − *j*| = 1 (except for K562 that, as discussed previously displays some anomalous behavior compared to the other cell lines). The data also show that *f*_*α*_ remains approximately constant for some range of genomic distances. This range appears to be somewhat cell-line dependent. As discussed in the main text the histogram analysis is more reliable for tag distances which are not too distant (|*i*− *j*| ≲ 5) as the differences between *R*_*α*_ and *R*_*β*_ are larger.

